# Phosphorylation of GMFγ by c-Abl coordinates lamellipodial and focal adhesion dynamics

**DOI:** 10.1101/248914

**Authors:** Brennan D. Gerlach, Guoning Liao, Kate Tubbesing, Alyssa C. Rezey, Ruping Wang, Margarida Barroso, Dale D. Tang

## Abstract

During cell migration a critical interdependence between *protrusion* and *focal adhesion* dynamics is established and tightly regulated through signaling cascades. Here we demonstrate that c-Abl, a non-receptor tyrosine kinase, can control these migratory structures through the regulation of two actin-associated proteins, glia maturation factor-_γ_ (GMF_γ_) and Neural Wiskott-Aldrich syndrome protein (N-WASP). Phosphorylation of GMF_γ_ at tyrosine-104 by c-Abl directs activated N-WASP (pY256) to the leading edge, where it can promote protrusion extension. Non-phosphorylated GMF_γ_ guides N-WASP (pY256) to maturing focal adhesions to enhance further growth. Antagonizing this signaling pathway through knockdown or mutation of tyrosine-104 to its non-phosphorylated form attenuates migration, whereas the phospho-mimic mutant GMF_γ_ enhances migration, thus demonstrating c-Abl, GMF_γ_, and activated N-WASP (pY256) as a critical signaling cascade for regulating migration in a primary human cell line.

---

Smooth muscle cell migration is critical for the development of the airways, vasculature, gut, and bladder^1-8^; however, it is also associated with the progression of pathologies, such as asthma, COPD, atherosclerosis, and vascular restenosis^9-13^. During migration, cells extend lamellipodium or protrusions, which are sheet-like extensions of plasma membrane that form through rounds of actin polymerization (assembly) and depolymerization (disassembly)^14,15,16,17^. Subsequently, signaling events trigger the connection of retrograde actin filaments to focal adhesion complexes to create a “molecular clutch”^18-21^. Focal adhesion complexes are nano-domain structures that attach cells to their extracellular matrix through the direct binding of heterodimeric transmembrane proteins called integrins^18,20^. Once the “molecular clutch” is formed, force generated by actin polymerization is transmitted to the ECM, resulting in traction force on the ECM and a net protrusion. However, the signaling mechanisms, that coordinate these events remain to be elucidated.

Glia Maturation factor-γ (GMF_γ_) (17kDa) is a member of the ADF/cofilin depolymerizing factor superfamily^22-30^. GMF_γ_ is expressed in a variety cell types^23,25,26,29,30^, whereas its homolog GMFβ is highly expressed in the brain^25,30^. Unlike its relative cofilin, GMF_γ_ does not interact directly with actin filaments, but rather binds specifically to Arp2 of the Arp2/3 complex to initiate actin branch disassembly^22,30^. The Arp2/3 complex has long been studied and is critical for the formation of lamellipodia^31,32,33^. Actin branch assembly is initiated through the interaction between Arp2/3 complex and nucleation-promoting factors such as Neural Wiskott-Aldrich syndrome protein (N-WASP) ^34-36^. Both Arp2/3 and N-WASP have been implicated in the formation of nascent focal adhesions within the lamellipodia^37-42^. Our previous reports found c-Abl, a non-receptor tyrosine kinase to be a direct regulator of GMF_γ_ through its phosphorylation at tyrosine 104 ^26^. Phosphorylation of GMF_γ_ decreased its binding affinity for Arp2, thus regulating actin polymerization and overall force generation in smooth muscle cells^26^. However, the function of GMF_γ_ phosphorylation and its influences on both lamellipodia and focal adhesions during cell migration remain to be elucidated.

Here we demonstrate in human airway smooth muscle cells (HASMCs), how the phosphorylation of GMF_γ_ at tyrosine 104 by c-Abl is a critical determinant for the spatial distribution of activated N-WASP (pY256) to the leading edge and to growing focal adhesions. Phosphorylated GMF_γ_ and N-WASP (pY256) were enriched at the leading edge, where it promoted extension of the lamellipodia. Conversely, expression of a non-phosphorylated mutant GMF_γ_ increased the enrichment of N-WASP (pY256) in focal adhesions where it increased growth. Altogether, our findings reveal c-Abl as a key regulator of migration through coordinating lamellipodial and focal adhesion dynamics.

## Results

### GMF_γ_ phosphorylation at Y104 regulates smooth muscle migration

To interrogate the function of GMF_γ_, we evaluated the effects of GMF_γ_ knockdown and rescue on smooth muscle cell migration. We generated stable GMF_γ_ knockdown HASMCs by using lentiviral particles encoding control or GMF_γ_ shRNA. Immunoblot analysis verified effective knockdown in cells expressing GMF_γ_ shRNA by 80% (Supplementary Fig. 1a). We performed a wound-healing assay to assess the effects of GMF_γ_ knockdown on directed cell migration. Loss of GMF_γ_ led to a decrease in the ability of cells to close the scratch area in 12hrs as compared to control shRNA expressing cells (Supplementary Fig. 1b-d).

Since our previous publication^26^ suggested that GMF_γ_ is important for actin re-organization in contraction, we wanted to elucidate the role of GMF_γ_ phosphorylation at this residue in cell migration. We engineered EGFP-tagged wild-type (WT) GMF_γ_, non-phosphorylated mutant (Y104F) GMF_γ_ (substitution of phenylalanine at Y104) and phosphorylation mimic mutant (Y104D) GMF_γ_ (aspartic acid substitution at Y104) (Supplementary Fig. 1e,f). These DNA constructs were transiently transfected into GMF_γ_ knockdown cells and plated onto 6-well collagen I coated dishes. Migration of live-cells was monitored by time-lapse microscopy and migratory parameters were analyzed using ImageJ software (Fig. 1a-i, Supplementary Video 1). Knockdown of GMF_γ_ led to decreases in speed, accumulated distance, and Euclidean distance, as compared to control shRNA expressing cells (Fig. 1a,b,f-i), which was consistent with the results evaluated by the wound scratch assay. Re-expression of WT-GMF_γ_ and Y104D-GMF_γ_ were capable of rescuing all migratory parameters associated with the knockdown effects of GMF_γ_ back to or greater than control shRNA expressing cells (Fig. 1c,e,f-i). Conversely, expression of Y104F-GMF_γ_ led to decreases in speed, directionality, accumulated distance, and Euclidean distance when compared to all other treatments (Fig. 1d,f-i). These findings suggest that GMF_γ_ is necessary for cell migration and its phosphorylation at Y104 regulates cell motility.

**Figure 1:**
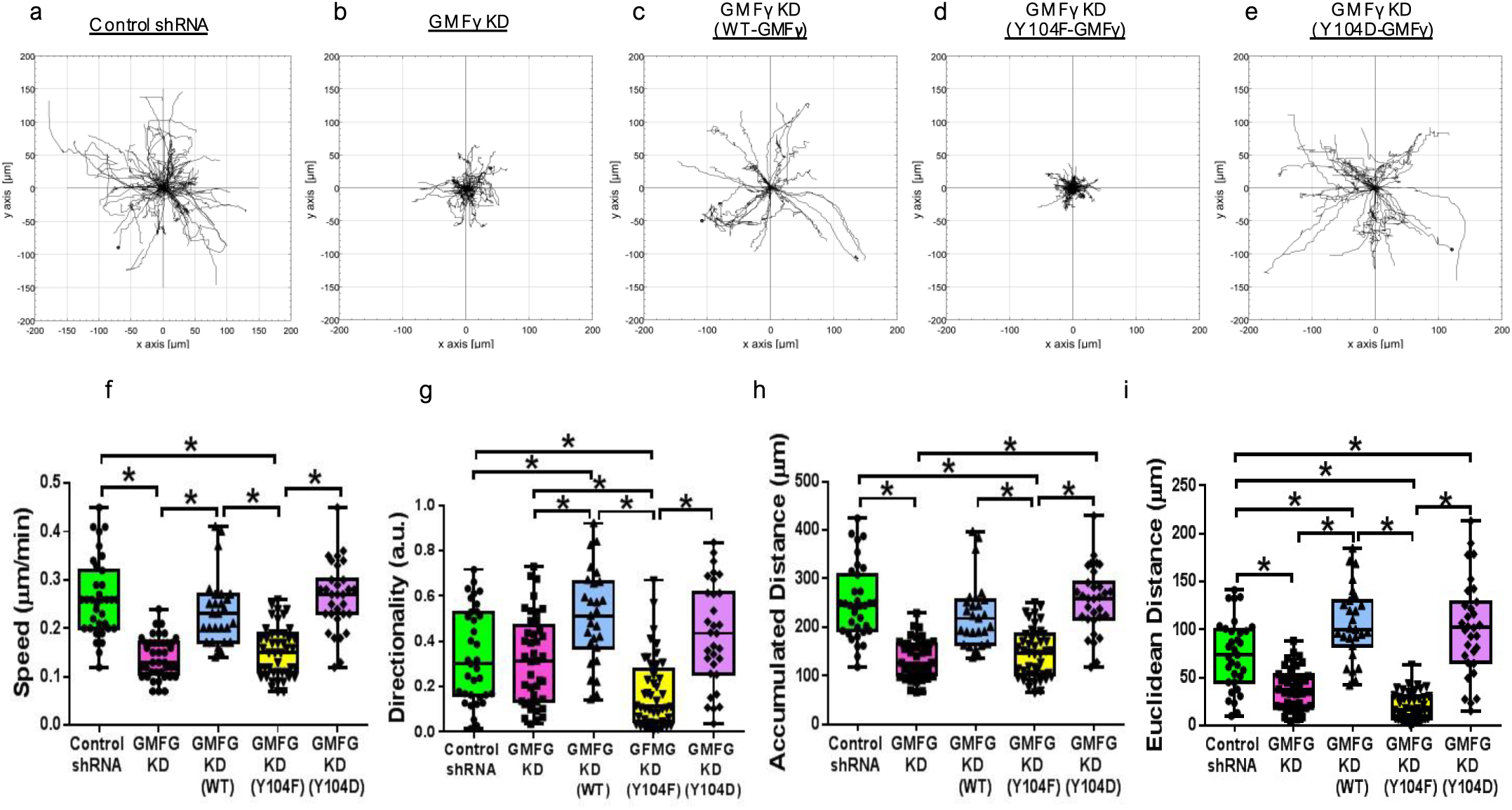
WT-GMF_γ_ and Y104D-GMF_γ_ expression rescues the attenuated migratory phenotype associated with knockdown of GMF_γ_. **(a-e)**. Time-lapse microscopy was used to track control shRNA and GMF_γ_ shRNA expressing cells, as well as rescue cells transfected using FUGENE HD (Promega): WT-GMF_γ_, Y104F-GMF_γ_, and Y104D-GMF_γ_ for 16 hours. Chemotaxis plots display migration patterns for each cell type. **Note: GMF_γ_ KD and Y104F-GMF_γ_ expressing cells display attenuated migration patterns as compared with Control shRNA, and re-expression of WT and Y104D-GMF_γ_. (f-i)**. Graphical comparisons represent the calculated speed (F(4, 174)=34.51), directionality (F(4, 174)=14.72), accumulated distance (F(4, 174)=33.81), and Euclidean distance (F(4, 172)=48.79) for each cell type. **Note: WT-GMF_γ_ and Y104D re-expression in knockdown cells rescued each migratory parameter back to control levels. Moreover, GMF_γ_ knockdown and re-expression of Y104F-GMF_γ_ were unable to rescue the migratory phenotypes**. Two-tailed one-way ANOVA with Tukey’s post hoc test was used (p<0.05, n=51, n=41, n=27, n=45, n=31).

### GMF_γ_ phosphorylation at Y104 promotes lamellipodial dynamics and F-actin formation

During migration, cells undergo the cyclic extension and retraction of lamellipodia to facilitate cell movement forward^15-17^. Our previous studies demonstrated that c-Abl can regulate smooth muscle migration, contraction, and proliferation through manipulating the actin cytoskeleton^9,26,46-51^. The finding that Y104D-GMF_γ_ rescued and enhanced directionality, an indicator of lamellipodial dynamics^15^, during migration spurred us to examine the role of this phosphorylation in lamellipodial dynamics.

To assess lamellipodial dynamics, smooth muscle cells were co-transfected with plasmids encoding EGFP-tagged recombinant GMF_γ_ and LifeAct-RFP plasmid, which generates a 17 amino-acid peptide to visualize F-actin in live cells^52^. Live-cell confocal and TIRF microscopy of cells expressing WT-GMF_γ_ revealed that the majority localized to both the near membrane region of the ventral side of the cell, and within lamellipodia and filopodia (Supplementary Fig. 1g and Supplementary Video 2). WT-GMF_γ_ was dynamically motile within the protrusion, where it moved to the leading edge and subsequently retreated during retraction of the lamella (Supplementary Video 2). Cells expressing WT-GMF_γ_ and Y104D-GMF_γ_ formed larger nascent protrusions compared to cells expressing Y104F-GMF_γ_ (Fig. 2a-c). In addition, cells expressing WT-GMF_γ_ and Y104D-GMF_γ_ displayed visible actin network in the protrusion, whereas the expression of Y104F-GMF_γ_ abolished actin network appearance in cells (Fig. 2a-c, and Supplementary Video 3a-c and Supplementary Video 4a-c).

**Figure 2:**
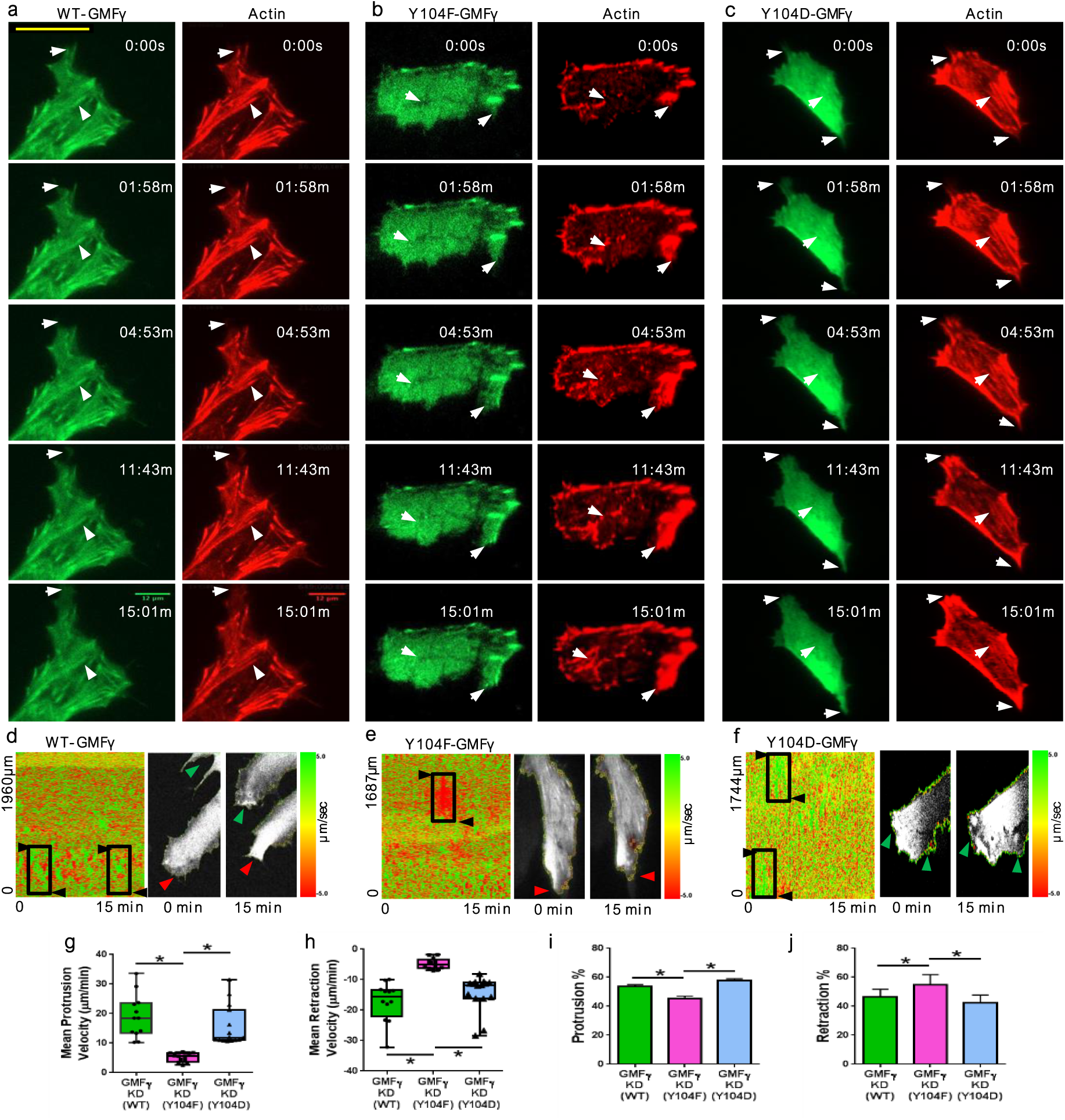
Y104F-GMF_γ_ disrupts lamellipodial dynamics and reduces the actin stress fiber network compared to WT-GMF_γ_ and Y104D-GMF_γ_. **(a-c)**. HASMCs were co-transfected with Life-Act RFP and EGFP-tagged WT-GMF_γ_, Y104F-GMF_γ_, or Y104D-GMF_γ_. GMF_γ_ and Actin localization was monitored every 1.3s for 15mins using the Zeiss Laser TIRF3 microscope and Zeiss LSM 880 confocal microscope with airyscan. **(d-e)**. Analysis of protrusion and retraction velocity was done using the ImageJ plugin ADAPT, which quantitates the protrusion boundary changes based on fluorescent intensity over time using a Triangle algorithm. **Note: the heat map displays higher velocities in green and lower velocities in red. Arrows indicate changes in lamellipodial positioning. Y104F-GMF_γ_ displays less protrusion and retraction velocities as compared to WT-GMF_γ_ and Y104D-GMF_γ_. (g,h)**. ADAPT quantifies the mean protrusion (F(2, 36)=18.05) and mean retraction (F(2, 36)=19.34) velocities compared between WT-GMF_γ_, Y104F-GMF_γ_, and Y104D-GMF_γ_. One-way ANOVA was used p<0.05, n=12, n=12, n=15. **(i,j)**. ADAPT also quantitates the percent protrusions (F(2, 36)=15.69) and retractions (F(2, 36)=15.75) observed during the time course. One-way ANOVA used p<0.05, n=12, n=12, n=15.

To analyze the lamellipodial dynamics, we used an ImageJ plugin known as “automated detection and analysis of protrusions” (ADAPT) (*Methods*) ^53^. Velocity heat maps displayed uniform protrusion/retraction velocities when cells expressed the WT-GMF_γ_ and Y104D-GMF_γ_ (Fig. 2d,f). However, larger areas of decreased velocities indicated in red were shown when cells expressed the Y104F-GMF_γ_ mutant (Fig. 2e). Expression of Y104F-GMF_γ_ severely impaired both protrusion velocity, as well as retraction velocity as compared to cells expressing WT-GMF_γ_ and Y104D-GMF_γ_ (Fig. 2g,h). Furthermore, expression of WT-GMF_γ_ and Y104D-GMF_γ_ exhibited 55% and 60% respectively more protruding lamellipodia, as compared to cells expressing Y104F-GMF_γ_ (Fig. 2i). On the other hand, 55% more retracting lamellipodia were observed with Y104F-GMF_γ_ expression compared to both WT-GMF_γ_ and Y104D-GMF_γ_ cells (Fig. 2j), suggesting a decrease in protrusion persistence^54^.These changes in lamellipodial dynamics suggest GMF_γ_ phosphorylation at Y104 is a critical site for regulating leading edge F-actin formation and extension of the leading edge, which could account for the improved directionality observed during migration of Y104D-GMF_γ_ expressing cells.

### Knockdown of GMF_γ_ disrupts N-WASP (pY256) spatial distribution and attenuates focal adhesion growth

Previous research has demonstrated that activated N-WASP (pY256) can regulate the Arp2/3 complex and promote both nascent focal adhesion and lamellipodia formation^38-42^. Since GMF_γ_ is known to regulate Arp2/3, we wanted to determine if it also regulated activated N-WASP at both the leading edge of the lamellipodium and at focal adhesions.

We first established an imaging assay where we only selected cells with prominent lamellipodia extending away from the nucleus. We captured images focused on the leading edge and focal adhesions lying within the lamellipodium and lamella respectively (Supplemental Fig. 2a, 3a). Immunostaining of the leading edge of HASMCs revealed colocalization of GMF_γ_, activated N-WASP (pY256), and Arp2 as assessed by Pearson’s correlation coefficient (Supplementary Fig. 2b,d). To determine whether endogenous phosphorylated GMF_γ_ also localized at the leading edge, we utilized a custom-made antibody targeting phosphorylated tyrosine 104^26^. Image analysis revealed that phospho-GMF_γ_ was enriched at the leading edge, but did not colocalize with Arp2, which is consistent with our previous studies^26^ (Supplementary Fig. 2c,d).

We next sought to determine if GMF_γ_ localized within focal adhesions by imaging the lamella (Supplementary Fig. 3a). HASMC were plated on collagen-coated coverslips and immunostained for GMF_γ_, vinculin (a marker for focal adhesions^18-22^), and zyxin (a marker for mature focal adhesions^43,44^. Confocal z-slice revealed that GMF_γ_ colocalized with both vinculin and zyxin (Supplementary Fig. 3b). To quantify the localization of GMF_γ_ within growing focal adhesions, we utilized Imaris 8.4.1 software analysis. We generated 3D rendered GMF_γ_ spots (green), as well as 3D rendered vinculin (purple) and zyxin (red) surfaces. We then masked vinculin surfaces that were contacting only zyxin surfaces (cyan) (*Methods* and Supplementary Fig. 3c). Using the masked channels, we could separate GMF_γ_ spots into distinct populations based on localization. Because of conservative rendering methods to avoid over estimation of protein overlap we found that 67% of GMF_γ_ spots were not localized to focal adhesions, whereas 33% were localized within focal adhesions. We next broke down the 33% into the amount of GMF_γ_ spots localized to either vinculin alone (40%), vinculin within a vinculin/zyxin positive focal adhesion (13%), localized to both vinculin/zyxin (25%), and localized to zyxin alone (13%) (Supplementary Fig. 3d). We also had a small population localized on the border of focal adhesions (9%) giving us a total of 6 distinct populations of GMF_γ_ (Supplementary Fig. 3d). This is the first evidence to our knowledge a portion of GMF_γ_ localizing to distinct regions of growing focal adhesions.

We subsequently wanted to determine if N-WASP (pY256) also localized with GMF_γ_ within focal adhesions. A confocal z-stack slice (Supplementary Fig. 3e) as well as 3D projection (Imaris) (Supplementary Fig. 3f) revealed that both GMF_γ_ and N-WASP (pY256) were colocalized together with vinculin as analyzed by Pearson’s coefficient (Supplementary Fig. 31, Supplementary Video 5). Moreover, co-immunoprecipitation analysis was used to assess interaction of GMF_γ_ with vinculin. Vinculin, N-WASP (pY256), and Arp2 were found in GMF_γ_ immunoprecipitates. Conversely, GMF_γ_, N-WASP (pY256), and Arp2 were found in vinculin precipitates (Supplementary Fig. 3g,m). Confocal microscopy also revealed N-WASP (pY256) localizing with GMF_γ_ and zyxin (Supplementary Fig. 3h,l). Imaris analysis generated GMF_γ_ (green) and N-WASP (pY256) (purple) spots, as well as zyxin (red) surfaces (Supplementary Fig. 3i,j). We then calculated the percentage of N-WASP (pY256) spots localizing to either vinculin or zyxin surfaces and found that 23% of N-WASP (pY256) localized with vinculin, whereas 19% localized with zyxin (Supplementary Fig. 3k). This is further supported with our Pearson’s coefficient analysis that shows N-WASP (pY256) is localized more with vinculin than zyxin (Supplementary Fig. 3l). The other 58% of N-WASP (pY256) is not localized within focal adhesions. Our results show that both GMF_γ_ and N-WASP (pY256) localize to the same regions within focal adhesions at similar proportions.

Since, GMF_γ_ and N-WASP (Y256) localize to focal adhesions, and knockdown of GMF_γ_ led to attenuated migration; this prompted us to examine whether GMF_γ_ plays a role in focal adhesion growth. We stained control shRNA and GMF_γ_ KD cells for N-WASP (pY256) and vinculin to examine focal adhesion morphology. Control shRNA cells exhibited large vinculin focal adhesions, whereas GMF_γ_ KD cells displayed reduction in focal adhesion size within the lamella (Fig. 3a,b). Total number of N-WASP (pY256) spots and vinculin surfaces were not significantly different between control cells and GMF_γ_ KD cells (Fig. 3c,d). However, the area, volume, and bounding box lengths (*Methods*) of vinculin surfaces were reduced in the GMF_γ_ KD cells as compared to control cells (Fig. 3 e-h). Furthermore, analysis of spots closest to surfaces, as well as Pearson’s coefficient for colocalization, revealed significantly decreased colocalization of N-WASP (pY256) to vinculin in the GMF_γ_ KD cells as compared to control cells (Fig. 3i,j).

**Figure 3:**
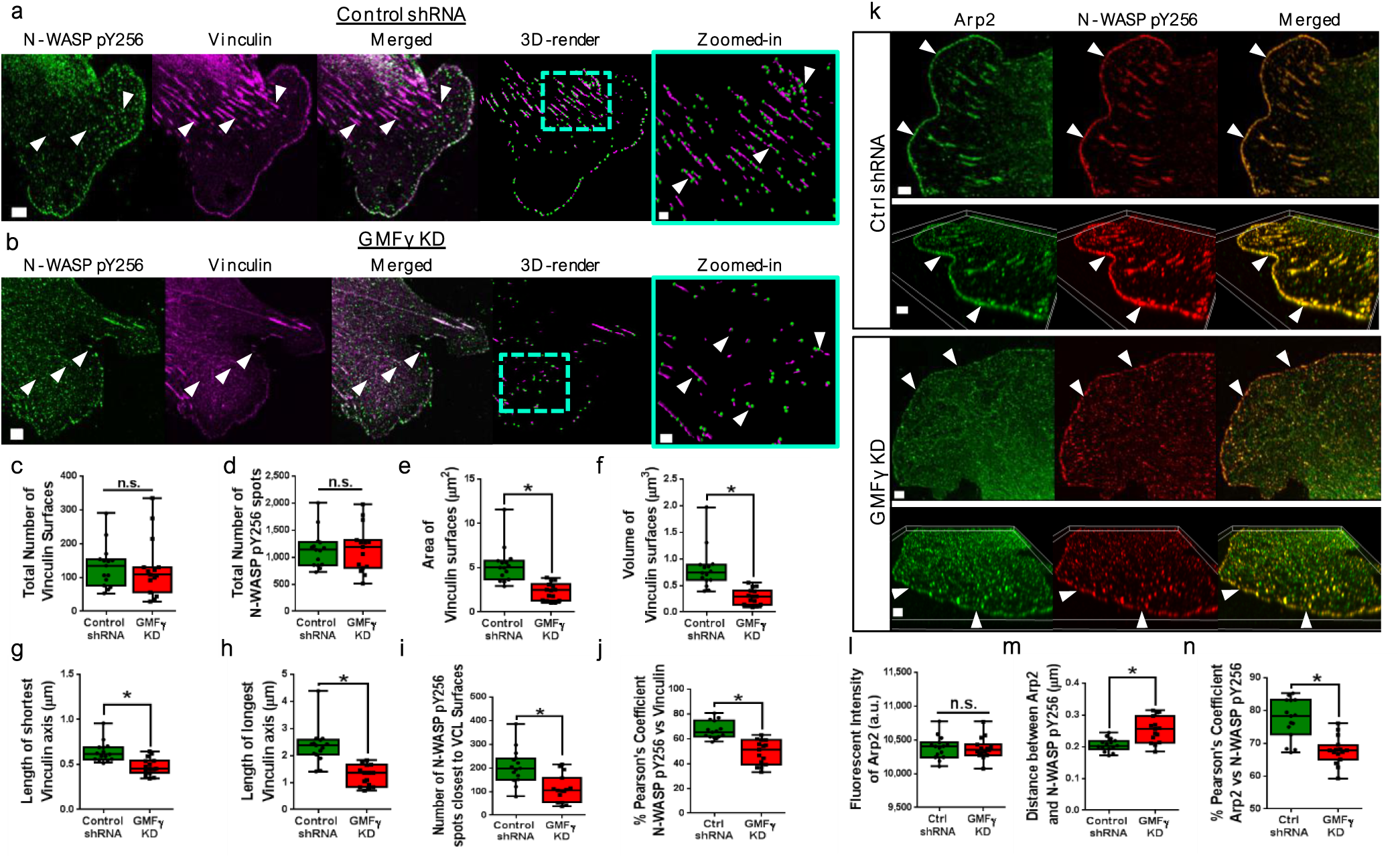
Knockdown of GMF_γ_ disrupts the localization of N-WASP (pY256) resulting in decreased focal adhesion growth. **(a,b)**. Control shRNA or GMF_γ_ KD cells were plated onto collagen coated coverslips and stained for N-WASP (pY256) (green) and Vinculin (magenta). Z-stack images were taken using the Zeiss 880 with Airyscan Scale bar 5μm, zoomed in images scale bar 1 urn. **(c-h)**. Imaris 8.4.1 software was used to generate 3D rendered spots and surfaces. Imaris software determined vinculin surface number (t=0.2417, df=28), N-WASP (pY256) spot number (t=0.07587, df=28), area (t=5.019, df=28), volume (t=4.813, df=28), length of shortest axis (t=3.673, df=28), length of longest axis (t=5.468, df=28). Students’ T-test was used p<0.05, n=15, n=15. **(i)**. Imaris determined the distance of N-WASP (pY256) spots to vinculin surfaces (t=3.828, df=28). Students’ T-test was used p<0.05, n=15, n=15. **(j)**. Imaris Coloc plugin was used to determine the Pearson’s coefficient for localization of N-WASP (pY256) and vinculin. Students’ T-test was used p<0.05, n=15, n=15, t=5.709, df=28. **(k)**. Control shRNA and GMF_γ_ KD were stained for Arp2 (green) and N-WASP (pY256) (red) Scale bar 5μm, 3μm. **(l)**. Imaris software was used to quantitate the fluorescent intensity of Arp2 between control shRNA and GMF_γ_ KD cells. Students’ T-test used with p<0.05, n=15, n=15, t=0.2182, df=28. **(m)**. Imaris software was used to generate 3D rendered to quantitate distance between Arp2 and N-WASP (pY256) spots. Students’T-test was used p<0.05, n=15, n=15, t=4.054, df=28. **(n)**. Pearson’s coefficient was determined using Imaris Coloc plugin for Arp2 and N-WASP (pY256) colocalization. Students’ T-test was used p<0.05, n=15, n=15, t=4.935, df=28.

We also checked the morphology and localization of Arp2 and N-WASP (pY256) at the leading edge in GMF_γ_ KD cells. Morphologically, GMF_γ_ KD cells displayed a less prominent leading edge as compared to control shRNA expressing cells (Fig. 3k). Control shRNA expressing cells displayed similar fluorescent intensity of Arp2 staining as GMF_γ_ KD cells, suggesting no changes in Arp2 expression (Fig. 3l). GMF_γ_ KD cells exhibited increased distance, as well as a decrease in Pearson’s coefficient from N-WASP (pY256) (Fig. 3m,n). Altogether these results suggest that GMF_γ_ can coordinate the positioning of N-WASP (pY256) to both focal adhesions and the leading edge.

### Y104F-GMF_γ_ mutant enhances focal adhesion growth and enrichment of N-WASP (pY256) to vinculin

We next postulated that the phosphorylation of GMF_γ_ may affect the spatial distribution of activated N-WASP, to the focal adhesions since we saw differential speeds of motile cells when Y104F-GMF_γ_ or YKMD-GMF_γ_ were expressed. To examine localization, we transfected GMF_γ_ KD cells with GFP-tagged WT-GMF_γ_, Y104F-GMF_γ_, or Y104D-GMF_γ_ and then immunostained for N-WASP (pY256) and vinculin. Imaging revealed WT-GMF_γ_ localized with N-WASP (pY256) in vinculin-positive focal adhesions and at the leading edge (Fig. 4a and Supplementary Fig. 4a). However, YKMF-GMF_γ_ and N-WASP (pY256) largely accumulated with vinculin while reducing their localization at the leading edge (Figure 4a and Supplementary Fig. 4a). Moreover, Y104D-GMF_γ_ and N-WASP (pY256) were largely absent within focal adhesions (Fig. 4a and Supplementary Fig. 4a). Expression of Y104F-GMF_γ_ significantly increased the number, area, volume, and bounding box lengths of vinculin-positive focal adhesions, as compared to both WT-GMF_γ_ and Y104D-GMF_γ_ (Fig. 4b-e). We then utilized the same 3D-rendered spot to surface masking strategy to establish the percentage of GMF_γ_ and N-WASP (pY256) spots localized with vinculin for each mutant-GMF_γ_ expressed. In cells expressing WT-GMF_γ_, a population of 72% GMF_γ_ and 68% N-WASP (pY256) were found to be outside (not contacting) vinculin, whereas a second population of 28% GMF_γ_ and 32% N-WASP (pY256) were contacting vinculin surfaces, which is consistent with our analysis of endogenous protein localization (Fig. 4f,g). In contrast, Y104F-GMF_γ_ expressing cells had a decrease in the percent of both GMF_γ_ and N-WASP (pY256) found outside of the focal adhesions to 46% and 55% respectively (Fig. 4f,g). This corresponded with an increase in GMF_γ_ (54%) and N-WASP (pY256) (45%) contacting vinculin surfaces, when Y104F-GMF_γ_ is expressed (Fig. 4f,g). Analysis of Y104D-GMF_γ_ expressing cells revealed a dramatic increase of GMF_γ_ and N-WASP (pY256) found outside of the focal adhesions with a percent of 89% and 86% respectively. This was followed by a decrease in both GMF_γ_ (11%) and N-WASP (pY256) (14%) found in focal adhesions (Fig. 4f,g). Altogether these results suggest that the phosphorylation state of GMF_γ_ is an important determinant for the localization of itself as well as N-WASP (pY256) to focal adhesions, which may influence their growth.

**Figure 4:**
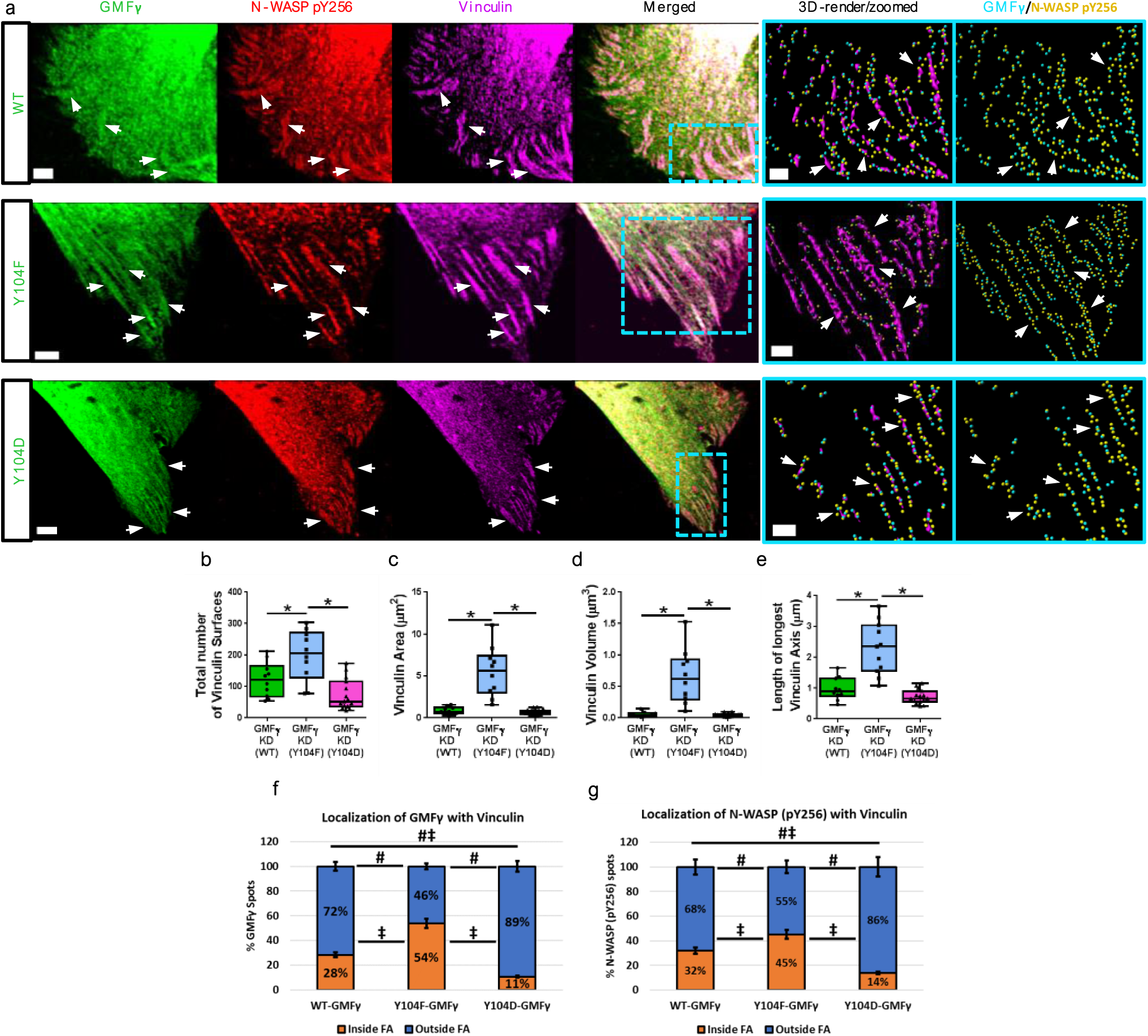
Expression of Y104F-GMF_γ_ increased the number and growth of focal adhesions through increased enrichment of N-WASP (pY256). **(a)**. GMF_γ_ KD cells were plated on collagen coated coverslips overnight and transfected with WT-GMF_γ_, Y104F-GMF_γ_, or Y104D-GMF_γ_ GFP tagged constructs. Cells were immunostained for N-WASP (pY256) and Vinculin. Z-stack images were taken using the Zeiss LSM 880 microscope with Airyscan scale bar 5μm. Arrows indicate focal adhesions localized with GMF_γ_ and N-WASP (pY256). Imaris software was used to generate 3D rendered spots and surfaces and gated on the spots closest to surfaces scale bar 1μm. **(b-e)**. Imaris software quantified the number (F(2, 32)=12.98), area (F(2, 32)=31.17), volume (F(2, 32)=25.08), and length of longest axis (F(2, 32)=39.25) of vinculin surfaces. **(f,g)**. Imaris was used to quantify the percentage of GMF_γ_ and N-WASP (pY256) spots localized outside or inside vinculin focal adhesions. Two-way ANOVATukey’s post hoc test used p<0.05, n=10, n=12, n =17.

We further wanted to investigate whether the expression of phosphorylated GMF_γ_ affected the formation of growing focal adhesions as identified by zyxin staining. GMF_γ_ KD cells were transfected with GFP-tagged WT-GMF_γ_, Y104F-GMF_γ_, or Y104D-GMF_γ_, then immunostained for zyxin and vinculin (Fig. 5a). Expression of Y104F-GMF_γ_ increased the number, area, volume, and bounding box lengths of vinculin focal adhesions, as compared to both WT-GMF_γ_ and Y104D-GMF_γ_ (Fig. 5b-e). Interestingly, the number of zyxin surfaces increased significantly in cells expressing Y104F-GMF_γ_ compared to both WT-GMF_γ_ and Y104D-GMF_γ_ cells (Fig. 5f). However, the area of zyxin surfaces did not change with expression of the GMF_γ_ constructs (Fig. 5g). Together this suggests that the phosphorylation state of GMF_γ_ may play a role in the growth of focal adhesions, which would explain the observed increase or decrease in migratory speed in cells expressing Y104D-GMF_γ_ or Y104F-GMF_γ_ respectively.

**Figure 5:**
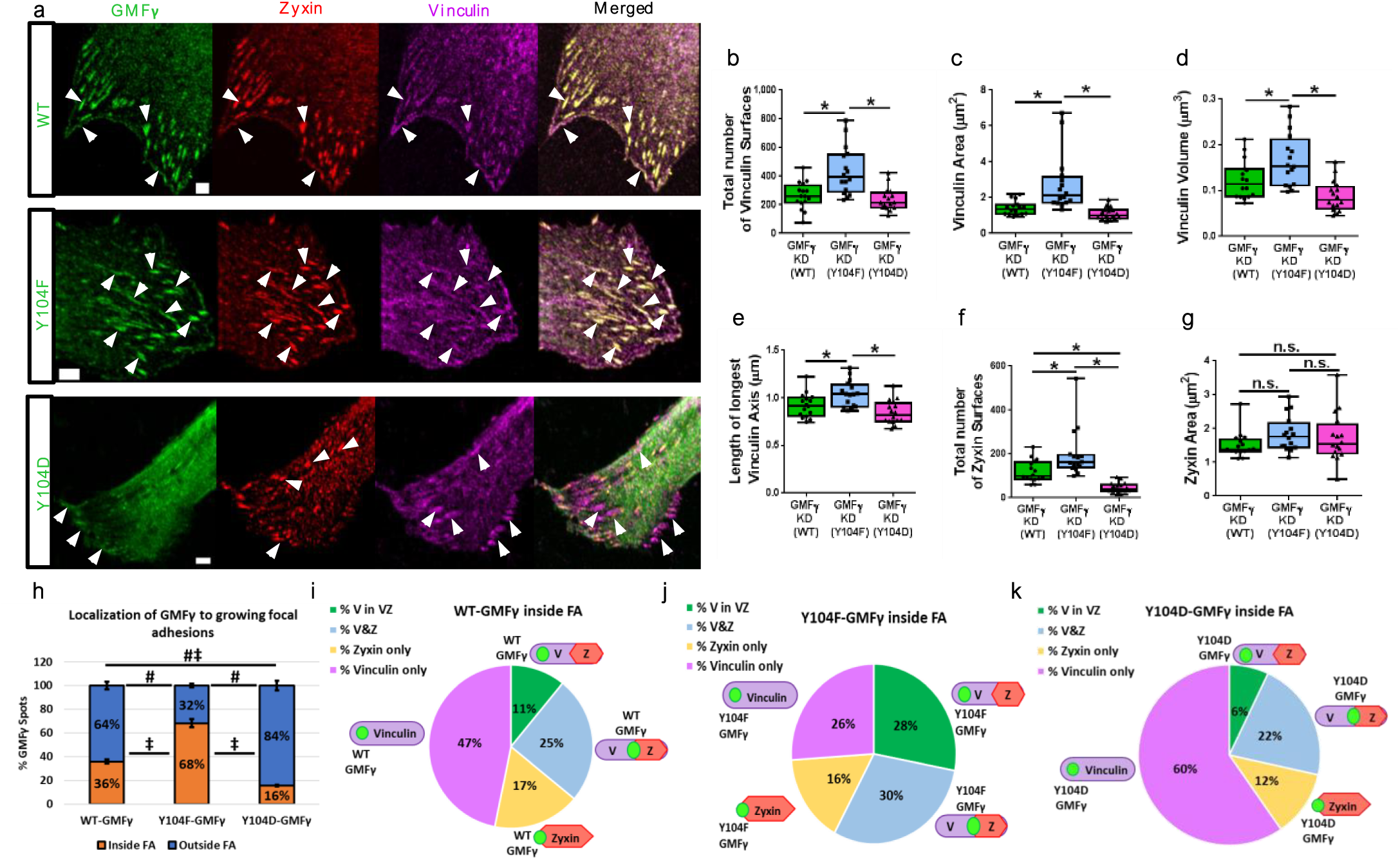
Mutation of Y104-GMF_γ_ modulates its regional distribution inside focal adhesions associated with variations in focal adhesion growth. **(a)**. GMF_γ_ KD cells were plated on collagen coated coverslips overnight and transfected with WT-GMF_γ_, Y104F-GMF_γ_, or Y104D-GMF_γ_ GFP tagged constructs. Cells were immunostained for zyxin and vinculin. Z-stack images were taken using the Zeiss LSM 880 microscope with Airyscan, scale bar 5μm. Arrows indicate localization of GMF_γ_, zyxin, and vinculin. Imaris software was used to generate 3D rendered spots and surfaces. **(b-e)**. Imaris software quantified the number (F(2, 45)=12.54), area (F(2, 45)=13.19), volume (F(2, 45)=12.02), and length of longest axis of vinculin surfaces (F(2, 45)=9.559). **(f,g)**. Imaris software quantified the number (F(2, 45)=18.40) and area of zyxin surfaces (F(2,45)=1.373). One-way ANOVA with Tukey’s post hoc test was used p<0.05, n=16, n=16, n=16. **(h)**. Imaris software was used to quantify the percentage of GMF_γ_ spots localized outside or inside focal adhesions. Two-way ANOVA with Tukey’s post hoc test was used p<0.05, n=16, n=16, n=16. **(i-k)**. Imaris software was used to mask vinculin surfaces contacting zyxin surfaces to quantify percent of GMF_γ_ spots within focal adhesions localized to vinculin only, zyxin only, vinculin in a vinculin/zyxin FA (%V in VZ), or both vinculin and zyxin (V&Z) for each GMF_γ_ mutant.

Additionally, we hypothesized that changes in GMF_γ_ phosphorylation may also change its localization within focal adhesions. We utilized our analysis protocol and discovered a consistent distribution of GMF_γ_ with increased localization inside focal adhesions when Y104F-GMF_γ_ (68%) was expressed as compared to WT-GMF_γ_ (36%) and Y104D-GMF_γ_ (16%) (Fig. 5h). Of the 36% of WT-GMF_γ_ localized within focal adhesions, the majority (47%) was localized with vinculin only, whereas 25% was localized with both vinculin and zyxin. The remaining 17% and 11% were localized to zyxin only and vinculin within a vinculin/zyxin focal adhesion respectively (Fig. 5i). Of the 68% of Y104F-GMF_γ_ localized to focal adhesions, the distribution was more uniform. Around 30% of Y104F-GMF_γ_ localized to both vinculin and zyxin, 28% was localized to vinculin within a vinculin/zyxin focal adhesion, 26% was localized to vinculin only and lastly 16% was localized with zyxin only (Fig. 5j). Conversely, of the 16% Y104D-GMF_γ_ localized to focal adhesions, the majority (60%) was localized to vinculin only. The rest of the distribution resembled that of WT-GMF_γ_ expressing cells with 22% localized to vinculin and zyxin, 12% zyxin only, and lastly 6% localized to vinculin within a vinculin/zyxin focal adhesion (Fig. 5k).

Ultimately, these results provide evidence that the localization of GMF_γ_ within different regions of the focal adhesion depends on its phosphorylation state and that this can increase the enrichment of active N-WASP within focal adhesions to increase growth.

### Expression of Y104D-GMF_γ_ rescues the migratory phenotype of c-Abl knockdown cells

c-Abl is critical in promoting migration of various cell types including smooth muscle cells^46,55-57^. Since c-Abl catalyzes the phosphorylation of GMF_γ_^26^, we questioned whether GMF_γ_ phosphorylation at Y104 mediates c-Abl regulated cell migration. We utilized time-lapse microscopy to evaluate the migratory parameters of control cells, c-Abl KD cells and KD cells expressing various GMF_γ_ constructs. Knockdown of c-Abl significantly decreased the migration of HASMC’s as compared to control shRNA expressing cells (Fig. 6a,b, f-i). Expression of WT-GMF_γ_ partially rescues the speed, accumulated distance, and Euclidean distance (Fig. 6c, f-i). However, expression of Y104F-GMF_γ_ did not rescue any of the migratory parameters (Fig. 6d, f-i). Furthermore, migratory parameters of cells expressing Y104D-GMF_γ_ were rescued to similar or greater than cells expressing control shRNA (Fig. 6e, f-i). These results suggest that the phosphorylation of GMF_γ_ at this residue is an important cellular switch that mediates c-Abl controlled cell migration.

**Figure 6:**
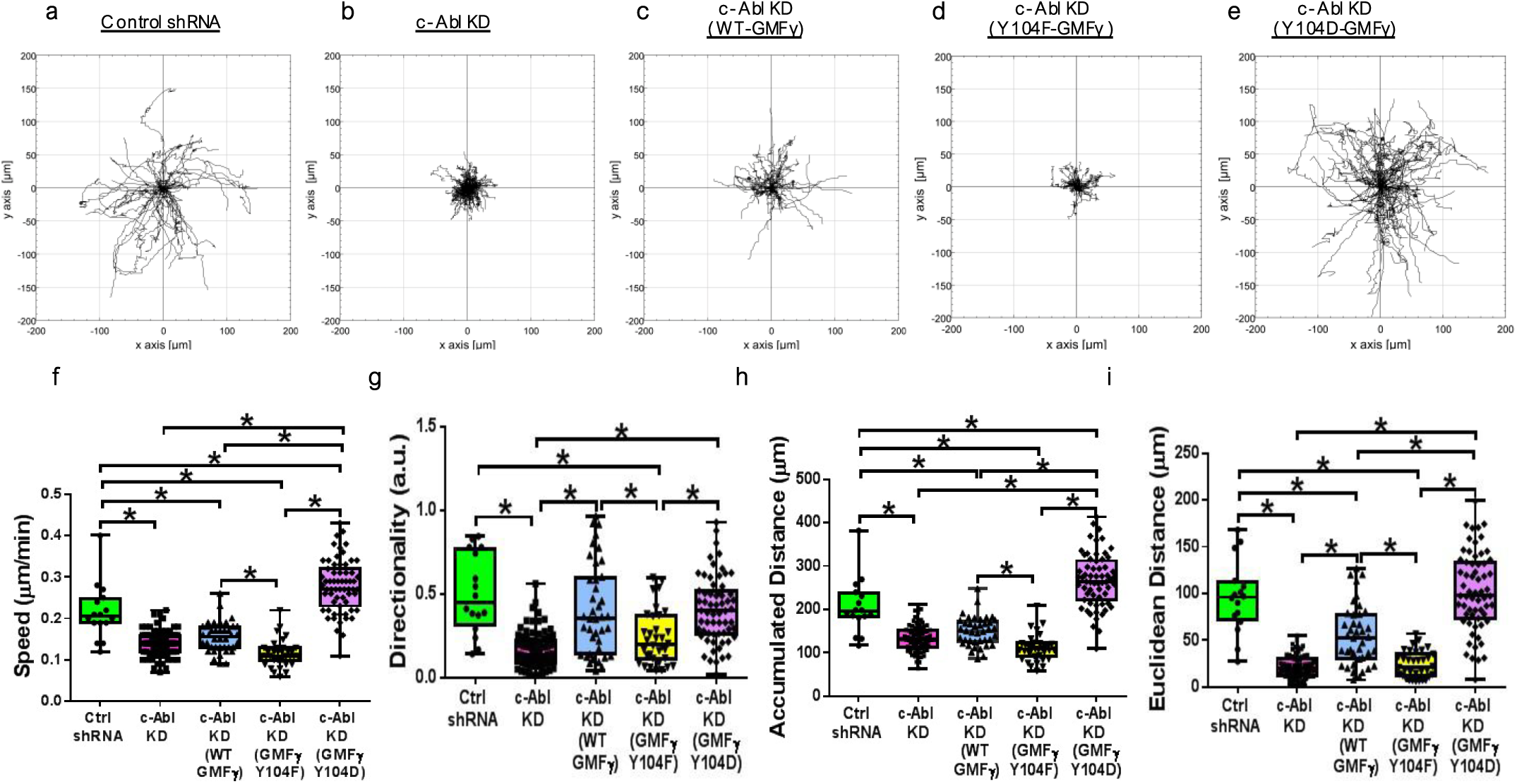
Re-expression of Y104D-GMF_γ_ rescues the migratory phenotype of c-Abl shRNA knockdown expressing HASMC. **(a-e)**. Time-lapse microscopy was used to track c-Abl lentiviral knockdown expressing cells, and the re-expression of WT-GMF_γ_, Y104F-GMF_γ_, and Y104D-GMF_γ_ constructs transfected using FUGENE HD (Promega) for 16 hours. Chemotaxis plots display migration patterns for each cell type. **(f-i)**. Graphical comparisons represent the calculated speed (F(4, 231)=112.1), directionality (F(4, 231)=24.71), accumulated distance (F(4, 231)=113.2), and Euclidean distance (F(4, 231)=88.73) for each cell type. One-way ANOVA with Tukey’s post hoc test was used (p<0.05, n=36, n=83, n=40, n=34, n=63).

### Knockdown of c-Abl enriches the spatial distribution of GMF_γ_ and N-WASP (pY256) to growing focal adhesions

To determine whether c-Abl affects the spatial distribution of GMF_γ_, cells expressing control shRNA and stable c-Abl KD cells were stained for GMF_γ_, N-WASP (pY256), vinculin and zyxin (Fig. 7a,b,i,j). Evaluation of c-Abl KD cells revealed an increase in vinculin area, volume, and bounding box lengths, but not in number when compared with control shRNA expressing cells (Fig. 7c-f). Similar to cells expressing Y104F-GMF_γ_, c-Abl KD cells also displayed increased enrichment of both GMF_γ_ and N-WASP (pY256) present in vinculin focal adhesions (Fig. 7g,h). Furthermore, we hypothesized that knockdown of c-Abl may increase the amount of mature focal adhesions explaining why these cells have attenuated migratory parameters. Assessment of zyxin staining in c-Abl KD cells exhibited an increase in zyxin number, but no change in area, volume, or bounding box lengths as compared to control shRNA expressing cells (Fig. 7k-n). This finding recapitulates the changes in focal adhesions observed in cells expressing Y104F-GMF_γ_. Moreover, c-Abl KD cells had greater enrichment of both GMF_γ_ and N-WASP (pY256) in zyxin containing focal adhesions (Fig. 7o,p). Overall these findings suggest that c-Abl coordinates focal adhesion growth through the regulation of GMF_γ_ and N-WASP (pY256) spatial distribution during cell migration.

**Figure 7:**
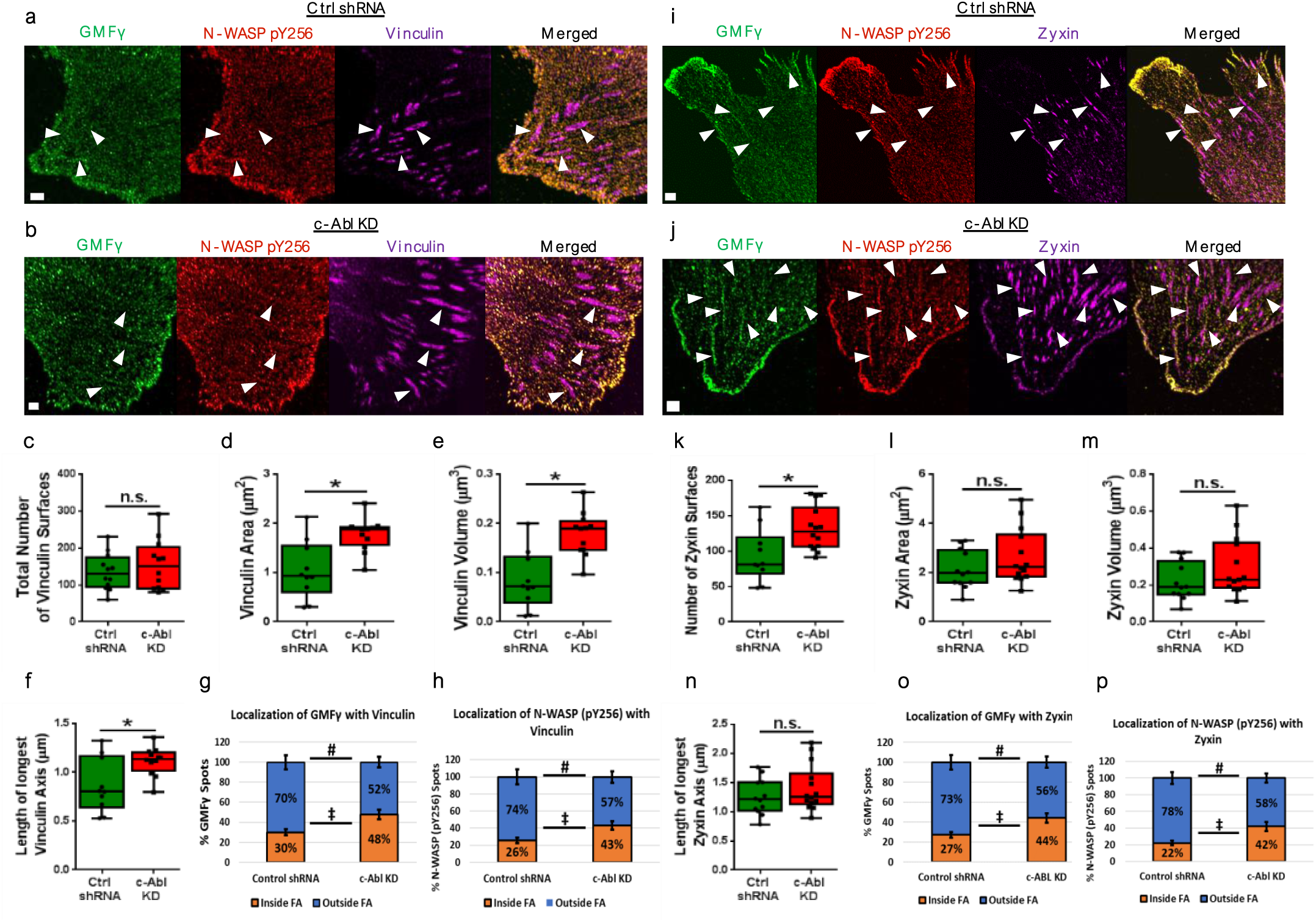
Knockdown of c-Abl increases focal adhesion growth, through enrichment of GMF_γ_ and N-WASP (pY256). **(a,b)**. Control shRNA and c-Abl KD expressing cells were plated on collagen coated coverslips for 60 minutes and immunostained for GMF_γ_, N-WASP (pY256), and vinculin. Images were taken using the Zeiss LSM 880 microscope with Airyscan, scale bar 5μm. Imaris software was used to generate 3D rendered spots and surfaces and gated on the spots closest to surfaces. **(c-h)**. Imaris software quantified the number (t=0.2724, df=20), area (t=3.670, df=20), volume (t=4.350, df=20), and length of longest vinculin axis. Imaris software was used to quantify the percentage of GMF_γ_ or N-WASP (pY256) spots localized outside or inside vinculin surfaces. Two-way ANOVA with a Tukey’s post hoc test was used p<0.05, n=14, n=17. **(i,j)**. Control shRNA and c-Abl KD expressing cells were plated on collagen coated coverslips for 60 minutes and immunostained for GMF_γ_, N-WASP (pY256), and zxyin. **(k-p)**. Imaris software quantified the number (t=2.748, df=22), area (t=1.927, df=22), volume (t=1.984, df=22), and length of longest zyxin axis (t=1.725, df=22). Imaris quantified the percentage of GMF_γ_ and N-WASP (pY256) spots localized outside or inside zyxin surfaces. Two-way ANOVA with a Tukey’s post hoc test was used (p<0.05, n=10, n=14).

## Discussion

Our results demonstrate that the phosphorylation of GMF_γ_ by c-Abl influences the spatial distribution of N-WASP (pY256) to coordinate lamellipodial and focal adhesion dynamics, important for directed cell migration (Fig. 8a). Here we show that the phosphorylation state of GMF_γ_ is a determinant for its localization, as well as activated N-WASP (pY256) to either the leading edge of the lamellipodia (Y104D-GMF_γ_) or to growing focal adhesions (Y104F-GMF_γ_) (Fig. 8a). Analysis of lamellipodial dynamics revealed that Y104D-GMF_γ_ increased recruitment of activated N-WASP to the leading edge of the lamellipodia to increase protrusion/retraction velocities. This is consistent with an observed increase in speed, directionality, and accumulated/net distance in migratory cells expressing Y104D-GMF_γ_. In contrast, Y104F-GMF_γ_ attenuated protrusion/retraction velocities and decreased the enrichment of N-WASP (pY256) at the leading edge. It is possible that Y104F-GMF_γ_ increases deb ranching of actin branches to promote actin retrograde flow and further actin re-organization within the lamella, thus slowing the extension of nascent protrusions (Fig. 8a,b). Whereas, enrichment of Y104D-GMF_γ_ at the leading edge may increase N-WASP (pY256) access to activate Arp2/3, leading to increased actin polymerization and subsequent protrusion extension (Fig. 8b). Still, synergy may exist with other depolymerizing factors within the lamella, such as cofilin, arpin, or coronin to drive actin retrograde flow and focal adhesion growth in place of GMF_γ_^58,59^. Live-cell super-resolution microscopy of WT-GMF_γ_ tagged with EGFP, demonstrated this dynamic movement of GMF_γ_ to the leading edge and subsequent retraction into the lamella, suggesting redistribution of a single pool of GMF_γ_. This is consistent with our image analysis where 67% of GMF_γ_ and N-WASP (pY256) were localized outside of focal adhesions, whereas 33% of GMF_γ_ and N-WASP (pY256) were localized inside the focal adhesions.

**Figure 8:**
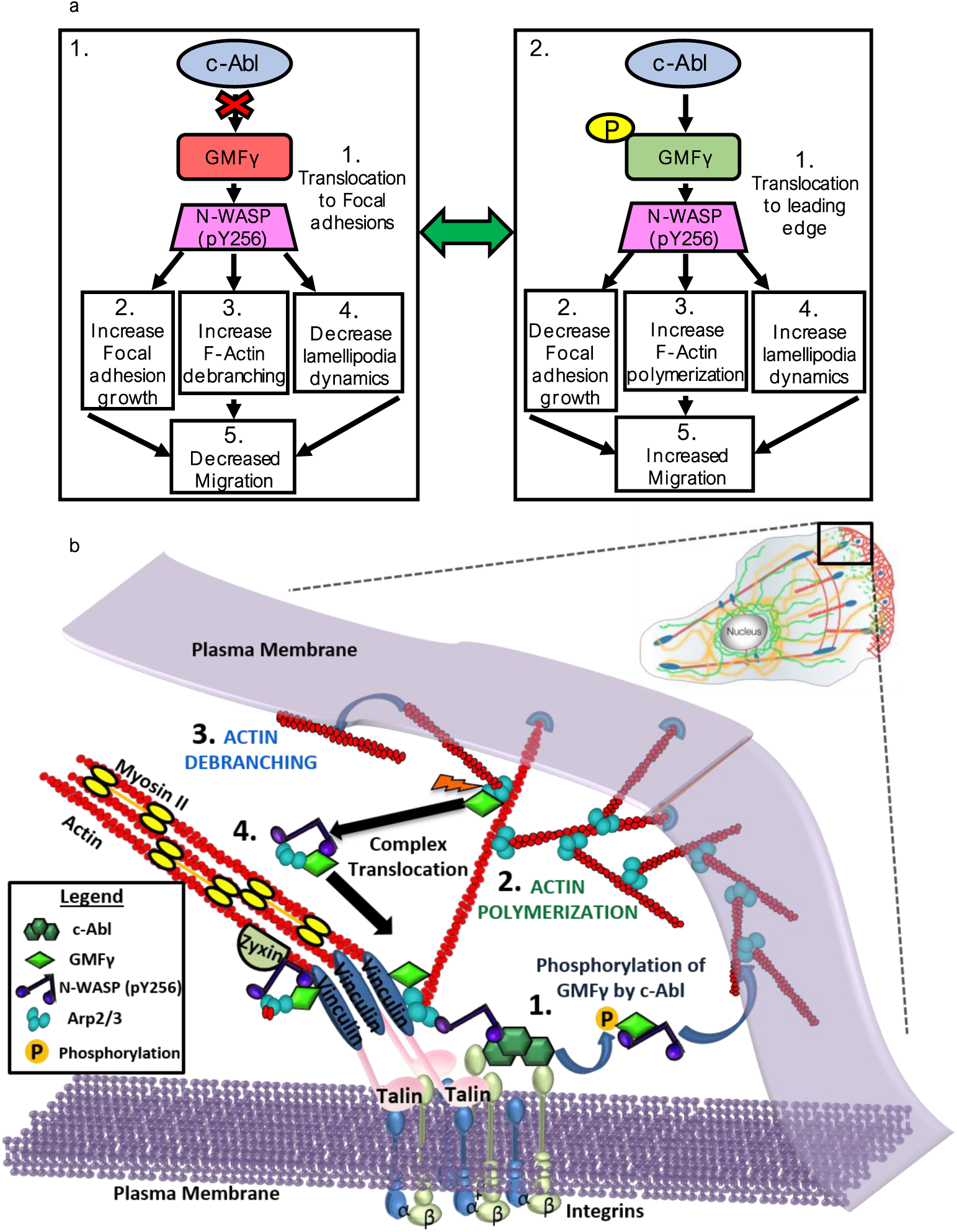
Model for the role of GMF_γ_ in the regulation of lamellipodial and focal adhesion dynamics during cell migration. This model is based on our results as described in greater detail within the Discussion. **(a). (1)** Non-phosphorylated GMF_γ_ guides N-WASP (pY256) to focal adhesions leading to increased growth. Non-phosphorylated GMF_γ_ also increased F-actin filament debranching, thus coupling focal adhesion growth with retrograde actin flow. This assists in engagement of the “molecular clutch” necessary for establishing force generation. Lack of GMF_γ_ and N-WASP at the leading edge causes decreased lamellipodial dyanmics. Too much non-phosphorylated GMF_γ_ can lead to decreased migration. **(2)** When c-Abl is activated, it phosphorylates GMF_γ_, which guides N-WASP (pY256) to the leading edge. This switch causes decreased N-WASP (pY256) enrichment within focal adhesions leading to limited growth. While at the leading edge active N-WASP can promote increased F-actin polymerization and promote extension of the lamellipodium causing increased migration. Altogether the coordination between non-phosphorylated and phosphorylated GMF_γ_ promotes cell migration. **(b)**. Interpreted model of how c-Abl signaling axis functions during migration. **(1)** Upon migratory stimulus, phosphorylation of GMF_γ_ by c-Abl leads to its translocation with N-WASP (pY256) to the leading edge, where N-WASP can activate the Arp2/3 complex to initiate actin polymerization **(2)** and protrusion formation. **(3)** Concomitantly, non-phosphorylated GMF_γ_ interacts with Arp2 within the lamella to drive actin branch disassembly, which promotes actin retrograde flow. **(4)** Translocation of the non-phosphorylated GMF_γ_, Arp2, N-WASP (pY256) complex to vinculin and zyxin further promotes the growth of focal adhesions to control migration. Collectively, the phosphorylation of GMF_γ_ and its translocation are critical in regulating both lamellipodial and focal adhesion dynamics, which is important for proper cell migration.

We further explored how GMF_γ_ and N-WASP (pY256) may influence the speed of migration through their localization to focal adhesions. Previous findings showed that focal adhesion growth rates are dependent upon F-actin flow, independent of vinculin^60^. Moreover, Arp2/3 was shown to interact directly with vinculin to promote focal adhesion formation and maturation^40^. Loss of GMF_γ_ caused decreased focal adhesion size and decreased enrichment of N-WASP (pY256), which may be due to lack of retrograde flow or lack of Arp2/3 activation within the focal adhesions respectively. On the other hand, it is likely that when GMF_γ_ is non-phosphorylated it can couple actin retrograde flow with focal adhesion growth by debranching Arp2/3 nucleated branches (Fig. 8a,b). Additionally, non-phosphorylated GMF_γ_ can distribute N-WASP (pY256) to the focal adhesion, which may promote growth through the activation of Arp2/3 complex, as an increase in focal adhesion number, area, volume, and length was observed with Y104F-GMFv expression (Fig. 8a,b). Increased focal adhesion growth coupled with increased actin flow has been shown to reduce traction forces^20,60^, ultimately leading to decreased speed of migration^20,60^, which we observed when Y104F-GMF_γ_ was expressed (Fig. 8a). We also observed a shift in localization of Y104F-GMFv towards focal adhesions containing both vinculin and zyxin rather than focal adhesions just containing vinculin. This suggests that Y104F-GMFv may target activated N-WASP to regions of maturing focal adhesions over more nascent adhesions, leading to increased engagement of F-actin with focal adhesions (Fig. 8b). In contrast, expression of Y104D-GMF_γ_ saw a decrease in enrichment within focal adhesions, which coincides with an accumulation of N-WASP (pY256) outside the focal adhesions and at the leading edge (Fig. 8a,b). Our previous findings showed that c-Abl localizes at the leading edge as well as within focal adhesions through direct interaction with β1-integrin^46^, providing spatial evidence of regulation at these two regions (Fig. 8b). Moreover, nascent adhesions are shown to form within the leading edge of the lamellipodia, whereas growing focal adhesions are enriched within the lamella^20,21,41^. Expression of Y104D-GMFv may limit the amount of activated N-WASP able to reach the growing focal adhesions within the lamella, which can limit the extent of growth, while also increasing its localization to the leading edge. There, enrichment of N-WASP (pY256) may activate the Arp2/3 complex to generate newly polymerizing actin branches necessary for protrusion extension (Fig. 8a,b). This may explain the significant rescue of speed and directionality observed with expression of Y104D-GMF_γ_ in migrating c-Abl KD cells. Clinically, reports have demonstrated that GMF_γ_, N-WASP, and c-Abl are all upregulated in several pathologies afflicted by enhanced cell migration, suggesting this signaling cascade may play a role^9, 39, 55, 61-64.^

In summary, our findings revealed a c-Abl, GMF_γ_, N-WASP (pY256) axis as an important signaling component for coordinating the interdependence between lamellipodial dynamics and focal adhesion growth to promote cell migration (Fig. 8b), which may have implications in other primary human cell types due to the ubiquitous expression of each protein.

## METHODS

Methods, including statements of data availability and any associated accession codes and references, are available in the online version of the paper.

## ACKNOWLEDGEMENTS

This work was supported by NHLBI grants HL-110951, HL-113208, and HL-130304 from the National Institutes of Health (to Dale D. Tang). Also, this work was supported by the American Heart Association predoctoral fellowship: 16PRE31430001 (to Brennan D. Gerlach). We acknowledge the Albany Medical College Imaging Core and Dr. J. Mazurkiewicz for helping us with the Zeiss LSM 880, Zeiss TIRF Laser3 and Imaris software.

## AUTHOR CONTRIBUTIONS

B.D.G performed and analyzed majority of experiments. G.L, A.R, and R.P. generated DNA constructs and shRNA knockdown cells. B.D.G., K.T., and M.B. performed imaging and Imaris analysis. B.D.G. wrote manuscript. B.D.G., K.T., M.B., and D.D.T. reviewed and edited manuscript. D.D.T. supervised the study.

## COMPETING FINANCIAL INTERESTS

The authors declare no competing financial interests.

## ONLINE METHODS

### Cell Culture

Primary Human airway smooth muscle cells were a generous gift from Dr. Reynold Panettieri Jr. (Rutgers, The State University of New Jersey, Rutgers Institute for Translational Medicine and Science, New Brunswick, NJ) and were maintained below passage 5 in Ham’s F12 media (ThermoFisher) with 10% fetal bovine serum (FBS, ThermoFisher) at 5% CO_2_ and 37°C. For experiments, cells were plated at 8,000 cells/ml onto collagen I coated coverslips (10μg/ml) for either 60 minutes or overnight (specified in figures). All live-cell experiments were performed using phenol-red free DMEM/F12 with HEPES, L-glutamine and contained 10% FBS (ThermoFisher, Gibco).

### Reagents and Transfection

cDNA encoding mTagRFP-T-Lifeact-7 (Addgene plasmid # 54586) was a gift from Michael Davidson (Florida State University, Tallahassee, FL). Human GMF_γ_ DNA (NCBI Accession Number, NM_004877.2) was synthesized by Life Technologies, and subcloned into bacterial vectors pEGFP. Quickchange II site-directed mutagenesis kit (Agilent Technologies, Santa Clara, CA) was used to generate Y104F mutant (phenylalanine substitution at Tyr-104). The 5’-primer was 5’-GCCGGAACAACAGATGATGTTCGCAGGGAGTAAAAACAGG-3’. The 3’-primer was 5’-CCTGTTTTTACTCCCTGCGAACATCATCTGTTGTTCCGGC-3’. Quickchange II site-directed mutagenesis kit (Agilent Technologies, Santa Clara, CA) was used to generate Y104D mutant (aspartic acid substitution at Tyr-104). The 5’-primer was 5’-GCCGGAACAACAGATGATGGATGCAGGGAGTAAAAACAG-3’. The 3’-primer was 5’-CTGTTTTTACTCCCTGCATCCATCATCTGTTGTTCCGGC-3’.

Primary HASM were transiently transfected with 1μg DNA using FuGene HD Transfection Reagent (Promega Corporation Madison, WI). Cells were transfected overnight and then media was changed to 10% FBS/F12. All transfected cell experiments were carried out 12-16hrs post-transfection.

Lenti-viral particles containing shRNA specific for GMF_γ_ or non-targeting control shRNA were purchased from Santa Cruz Biotechnology. HASM cells were infected with control shRNA lentivirus (Cat#sc-108080) or GMF-_γ_ shRNA lentivirus (Cat#sc-97348-V) for 12hrs. They were then cultured for 3-4 days. Positive clones expressing shRNAs were selected by puromycin. Immunoblot analysis was used to determine the expression levels of GMF-_γ_ in these cells. GMF-_γ_ KD cells and cells expressing control shRNA were stable at least five passages after initial infection. In addition, stable c-Abl KD cells were previously characterized by our laboratory^46,49,51^.

Primary antibodies used for immunofluorescence (IF) and western blot (WB): anti-goat total GMF_γ_ (E-13) (1:25_IF, 1:50_WB, Santa Cruz Biotech Lot# B0915, Cat# sc-168016), anti-rabbit N-WASP Y256 (1:25_IF, 1:100_WB, EMD Millipore Lot# 2795491, 2838736, Cat# AB1966), anti-mouse Arp2 (E-12) (1:25_IF Santa Cruz Biotech Lot# B0612, Cat# sc-166103), anti-mouse vinculin (1:25_IF Invitrogen, ThermoFisher Lot# RA 2147394, Cat# MA1-34629), anti-rabbit vinculin (1:25_IF, 1:100_WB Sigma Aldrich), anti-mouse zyxin (1:25_IF Santa Cruz Biotech Lot# B1317, Cat# sc-293448). The anti-rabbit phospho-GMF_γ_ (Tyr-104) antibody was custom made by Thermo Scientific (Pierce). The sequence of the peptide for generating phospho-GMF-γ antibody was CKPEQQMMY(P)AGSKNRLVQTA (NCBI Accession Number, NM_004877.2). Secondary antibodies were all purchased from Invitrogen (ThermoFisher), which includes Alexa 405, 488, 555, 546, and 647 at a concentration of 1:200.

### Western blot

Cells lysed with 2x SDS sample buffer composed of 1.5% dithiothreitol, 2% SDS, 80 mM Tris-HCl (pH 6.8), 10% glycerol and 0.01% bromophenol blue were boiled for 5mins and separated onto SDS PAGE, then electro-transferred to nitrocellulose paper. Membranes were blocked using 2% bovine serum albumin (BSA) in PBS for lhr and then probed with specific primary antibodies followed by horseradish peroxidase-conjugated secondary antibody (Fisher Scientific). Proteins were visualized with Amersham enhanced chemiluminescence (ECL) select Western Blotting Detection Reagent (GE Healthcare Lot# 9774291 Cat# 45000999PM) using the Amersham Imager 600 (GE Healthcare). The levels of total protein or phosphoprotein were quantified by scanning densitometry of immunoblots (IQTL software by GE Healthcare).

### Co-lmmunoprecipitation

Cells were plated in 100mm cell culture dishes and allowed to grow to 90-100% confluence. Media was changed to SFM overnight. Media was removed and Lysis buffer (10% Trition X 100, 0.5M EDTA, 1M HEPES pH 7.6, 20% SDS, 100x Halt protease/phosphatase single-use Inhibitor cocktail Cat# 78442 Lot# RH235957) was added and incubated at 4°C for 10 minutes. Cells were scraped into 1.5ml tubes and rotated for lhr at 4°C. Centrifugation at 13.0rpm (x1000g) for 20 minutes and removal of supernatent was placed in new 1.5ml tube. Addition of Protein A/G Plus-agarose (Santa Cruz Biotech Lot# B0116 Cat# sc-2003) rotated for 30 minutes at 4°C. Centrifuge for 5 minutes and removal of supernatent and placed in new 1.5ml tube. Subsequently, addition of 2μg primary antibody or 1.5(0.1 of normal IgG as control were added to supernatent and rotated overnight at 4°C. Following incubation, Protein A/G with agarose was added and rotated for 3 hours at 4°C then centrifuged for 15 minutes at 13.0 rpm (x1000g) and removal of supernatent. Pellets were resuspended in IP Wash Buffer (5M NaCl, 1M Tris pH 7.6, 10% Triton x100, and H_2_O), centrifuged and the supernatents discarded 4 times. Pellets were saved and mixed with 2x SDS buffer, then boiled for 5 minutes and spun down. Co-IP samples were loaded and separated on an SDS PAGE gel, where they were electro-transferred onto nitrocellulose paper and immunoblotted with the correct primary antibodies. Visualization of co-IP was carried out using Amersham ECL (GE Healthcare) and imaged using the Amersham Imager 600 (GE Healthcare).

### Immunofluorescence

Cells were plated onto collagen I coated coverslips for 60mins, then fixed using 4% paraformaldehyde for 15minutes at room temperature and then permeabilized with 0.2% Triton X 100 in PBS. Coverslips were washed 3x for 5minutes with lx PBS in between each step. Coverslips were blocked using 2% BSA/PBST for 30minutes then primary antibodies were added with 2% BSA/PBST and incubated for lhr each. Secondary antibodies were added at a concentration of 1:200 for lhr each. Coverslips were fixed onto slides using Prolong Diamond mounting medium (ThermoFisher).

### Microscopy

Super-resolution microscopy was used for fixed-cell and live-cell imaging conducted on a Zeiss LSM 880 NLO confocal microscope with Airyscan (Carl Zeiss Microscopy Jena, Germany) equipped with 63x oil 1.4 numerical aperture (NA) objective lens (Huff et. al, 2015, Huff, J. 2015, Huff, f. 2016). Microscope software used is the Zen Black 2 edition to process Super-resolution images for the Airyscan. Microscope has incubation chamber set to 30-37°C and has 5% CO_2_ for live-cell imaging. Lasers used in each fixed-cell experiment were Argon (488nm wavelength), Diode 405-30 (405nm), DPSS 561-10 (561nm), HeNe633 (633nm). Z-stacks were acquired at an average of 11 slices with 0.19|xm distance between each slice. Live-cell imaging of GMF_γ_-GFP tagged and Life-Act-RFP constructs, utilized the Fast-Airyscan module on the Zeiss LSM 880 confocal microscope. Z-stack live-images were taken at an average of 6 slices with 0.19mn distance between slices for 15 minutes at a frame rate of 15 seconds. A Zeiss TIRF Laser3 Microscope was used to achieve resolution at 200nm along the z-axis for analysis of GMF_γ_ constructs localization to vinculin-positive focal adhesions. A TIRF angle of-73.4° (64nm depth) was used along with the lasers 488nm, 405nm, and 565nm. Time-lapse microscopy was achieved by using a Leica A600 microscope with a 6-well incubator chamber hooked up to 5% CO2. Epifluorescence imaging was utilized for cells that were transfected with fluorescent protein constructs. Only cells expressing a specific fluorescent construct were imaged and used for analysis. Epifluorescence was achieved by using a GFP filter cube. Multi-position marking function was used to designate specific x and y coordinates for time-lapse capture of images every 10 minutes for a 16-hour experimental period.

### Image Analysis

#### Colocalization Analysis

Imaris Software 8.4.1 was used to analyze z-stack fixed-images of HASM stained for total GMF_γ_, phospho-GMF_γ_, Arp2, N-WASP Y256, Vinculin, and Zyxin. Imaris “Coloc” plugin was used to quantify pixel-by-pixel images thresholded for intensity to remove background noise. Pearson’s coefficient was used to determine the degree of localization between two channels at each pixel. Colocalization is determined by an r-value > 0.5 for an n=20 cells.

#### Spots to Surfaces Analysis

Imaris 8.4.1 was used to generate and analyze the distance between spots and surfaces. An algorithm was utilized to create surfaces based upon fluorescent intensity and volume. Once surfaces were created, Imaris plugins were used to measure the total number, area, volume, and bounding box lengths (length of shortest axis, length of second longest axis, and length of longest axis) of each surface. If the image contained two surfaces (vinculin and zyxin), a mask of each surface was created. Any voxel outside of the masked surface is signified as having a 0.00 intensity value. This allows separation of channels inside regions within a masked surface that is contacting both vinculin and zyxin or just vinculin alone. A separate algorithm was used to create spots based upon fluorescent intensity and quality for either GMF_γ_ or N-WASP Y256 channels. Once spots were created, a plugin was used to measure the total number of spots for specific channels. Also, a separate plugin was used to threshold on spots closest to surfaces. From there a percentage of the number of spots closest to surface to total number of spots could be generated for analysis.

#### Protrusion/Retraction Analysis

The Imagef plugin ADAPT (automated detection and analysis of protrusions), which utilizes the fluorescent intensities at the cell border to measure changes in protrusion/retraction position and velocity overtime, was used to analyze Zeiss LSM 880 confocal live-cell movies [53]. Input of the spatial resolution, frames per minute, thresholding method (Triangle), smoothing filter radius, erosion iterations, spatial filter radius, temporal filter radius, cortex depth, visualization line thickness, and minimum/maximum velocity parameters will generate protrusion boundaries based on changes in fluorescence intensity. Once program has analyzed the movie it will generate analysis based on protrusion trajectory, protrusion velocity, and retraction velocity. It will also generate a velocity heat map that can be used to visualize changes in lamellipodial dynamics.

### Statistical Analysis

All data was analyzed using GraphPad Prism version 6.00 software (Windows, GraphPad Software, La Jolla, CA). A two-tailed one-way ANOVA followed by Tukey’s multiple comparisons test was used for comparing Pearson’s correlation coefficient data generated from Imaris 8.4.1 colocalization analysis (significance was determined by a p<0.05). A two-tailed one-way ANOVA followed by Tukey’s multiple comparisons test was also using for comparing Time-lapse microscopy parameters (significance was determined by a p<0.05). For each ANOVA the F-value and degrees of freedom are denoted by F(df, df)=value. A two-tailed Student’s T-test was used to determine a significance of p<0.05 for focal adhesion parameters, colocalization between two channels, protein level expression, and “wound” closure rates. For each Student’s T-test the t-value and degrees of freedom are denoted as follows t=value, df=value. Box and whisker plots were used to represent the data shown.

